# Xenomake: a pipeline for processing and sorting xenograft reads from spatial transcriptomic experiments

**DOI:** 10.1101/2023.09.04.556109

**Authors:** Benjamin S Strope, Katherine E Pendleton, William Z Bowie, Gloria V Echeverria, Qian Zhu

## Abstract

Xenograft models are attractive models that mimic human tumor biology and permit one to perturb the tumor microenvironment and study its drug response. Spatially resolved transcriptomics (SRT) provide a powerful way to study the organization of xenograft models, but currently there is a lack of specialized pipeline for processing xenograft reads originated from SRT experiments. Xenomake is a standalone pipeline for the automated handling of spatial xenograft reads. Xenomake handles read processing, alignment, xenograft read sorting, quantification, and connects well with downstream spatial analysis packages. We additionally show that Xenomake can correctly assign organism specific reads, reduce sparsity of data by increasing gene counts, while maintaining biological relevance for studies.

## Introduction

Xenograft models, including patient derived xenografts (PDXs) and cell line xenografts, are a widely used component of cancer research for understanding tumor/stroma interactions, screening drug therapeutics, and simulating human tumor biology to understand cancer progression and therapy resistance [1–3]. With the rising popularity of spatially resolved transcriptomic technologies, there is a growing need for processing pipelines that can handle reads from PDX samples. Sequencing experiments from a PDX sample often contains a mixture of reads originating from both the host and graft genomes. A unique challenge is unambiguously assigning mRNA reads as belonging to host and graft transcriptomes [4] This problem is important to resolve because in order to develop into a viable xenograft, the host and graft organisms must exhibit a strong degree of homology [5], which often leads to ambiguous mapping of reads to either organism [4]. Currently, there are limited options available in standard spatial pipelines to handle reads from PDX samples.

Space Ranger, a popular spatial processing platform [6], recommends building an integrated reference assembly containing both human and mouse genes to which PDX reads are mapped using the aligner STAR [7]. Doing so treats homologous genes as if they are paralogs within the same organism, within which reads will map to the organism’s genomic location with the higher alignment score. Because of the high degree of homology between human and mouse, there are many locations in the human and mouse genomes where reads are aligned with almost equal probability. Another approach is to first align PDX reads to the graft genome and map the remaining unaligned reads to the host [8,9], however, the results of this approach will depend on the order of execution since the first organism being aligned to will receive most of the mappings. To address these issues, precise methods such as Xenome [10] and Xengsort [11] have been developed to enable a sensitive and alignment-free way to classify reads as belonging to the graft and host genomes. While these tools have so far worked on bulk samples, they have not been adapted to work on spatial sequencing data from 10X Genomics and Visium platforms [12,13]. Technically speaking, such an adaptation would require tools to properly recognize and handle cellular barcodes/ spatial barcodes and unique molecular identifier (UMI) [14] information from read sequences to further apportion reads to organisms and spatial locations. Current Xengsort and Xenome implementations lack this capability.

To facilitate the adoption of spatial transcriptomics for PDX studies, we thus have developed a pipeline, Xenomake, which is an end-to-end pipeline that includes read alignment and gene quantification steps for xenograft reads generated by spatial transcriptomic platforms and uses a xenograft sorting tool to apportion these reads to the host and graft genomes. Xenomake (https://github.com/istrope/Xenomake) is written based on Snakemake [15] and is fully open source. We evaluate Xenomake by conducting comparisons with the existing solution, Space Ranger [6], and show the superiority of our tool. Throughout, we demonstrate the application of Xenomake on a published medulloblastoma PDX spatial transcriptomic (ST) dataset [16] and a newly generated triple negative breast cancer (TNBC) PDX ST dataset.

## Results

### Xenomake workflow

Xenomake (https://github.com/istrope/Xenomake) is a xenograft reads sorting and processing pipeline adapted for spatial transcriptomic data. It consists of the following steps: read tagging/trimming, alignment, annotation of genomic features, xenograft read sorting, subsetting bam, filtering multi mapping reads, and gene quantifications (**Figure 1**). In the first step, spatial barcodes and UMI information are extracted from individual reads and tagged to the reads to generate an unaligned tagged BAM file. Then, the reads are independently aligned to human and mouse genomes using STAR sequence aligner. Reads are extracted where they are simultaneously aligned to both genomes, forming an ambiguously mapped reads list (also called species-overlapping reads). These overlapping reads are subject to Xengsort K-mer alignment-free method to classify reads as belonging to host, graft, both, ambiguous, or neither category. Each category forms an inclusion reads list which we use to subset the original STAR-aligned bam file. The final step of the pipeline performs read multimapping handling and gene expression quantification (**Figure 1**).

**Figure 1:**
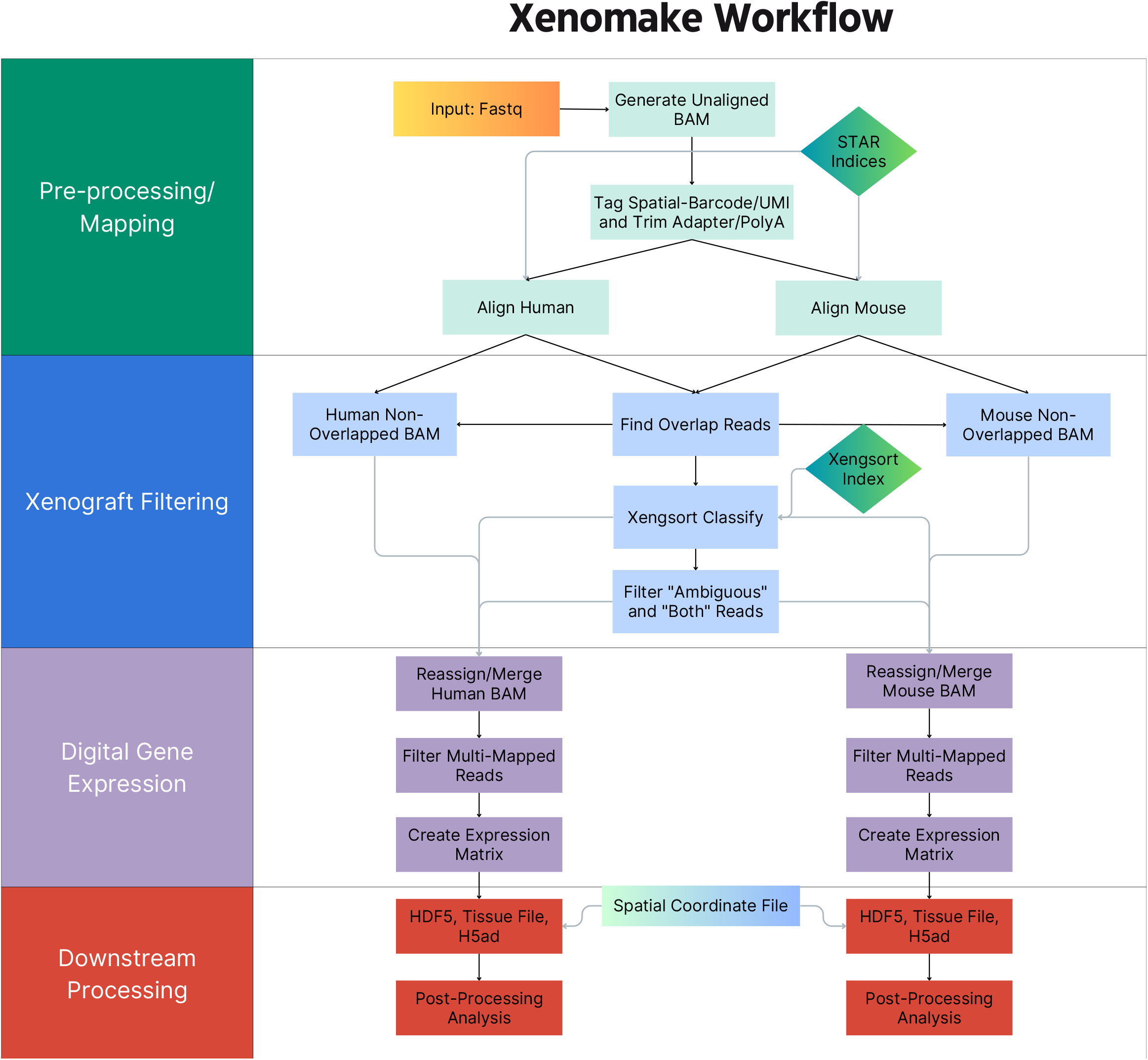
Overview of the Xenomake Pipeline. The graph shows all the processing, mapping, xenograft sorting, expression, and downstream processing performed by Xenomake. Arrows show the order of rules performed starting with a Fastq input.

Notably, Xenomake contains a policy regarding handling reads classified as both and ambiguous by Xengsort. Xenomake differs from others in that it adopts a flexible strategy to resolve both/ambiguous categories to make reads in these categories usable, rather than removing them. By default, Xenomake uses the genomic location (exonic, intronic, intergenic, or pseudogene) to determine the best aligned location of a multimapping read. A multimapper favors the exonic alignment over intergenic, pseudogenic, and any other secondary alignments. Otherwise, if both organisms produce exonic alignments, Xenomake uses the alignment score to further assign the read to the higher scored organism. The output of the pipeline is a barcode-by-gene expression matrix which can be in either Comma-Separated Values (CSV) or Hierarchical Data Format version 5 (HDF5) format [19], making it fully compatible with downstream spatial tools such as ScanPy [20], Seurat [21], and Giotto [22].

### Xenomake increases gene counts and reduces sparsity of spot-by-gene matrices

To illustrate the capability of Xenomake, we generated a ST dataset for a previously characterized TNBC PDX model PIM001P [9,23] generated from a treatment naive TNBC patient who went on to exhibit therapeutic resistance and aggressive disease progression (see Methods) (**Figure 2a**).

**Figure 2:**
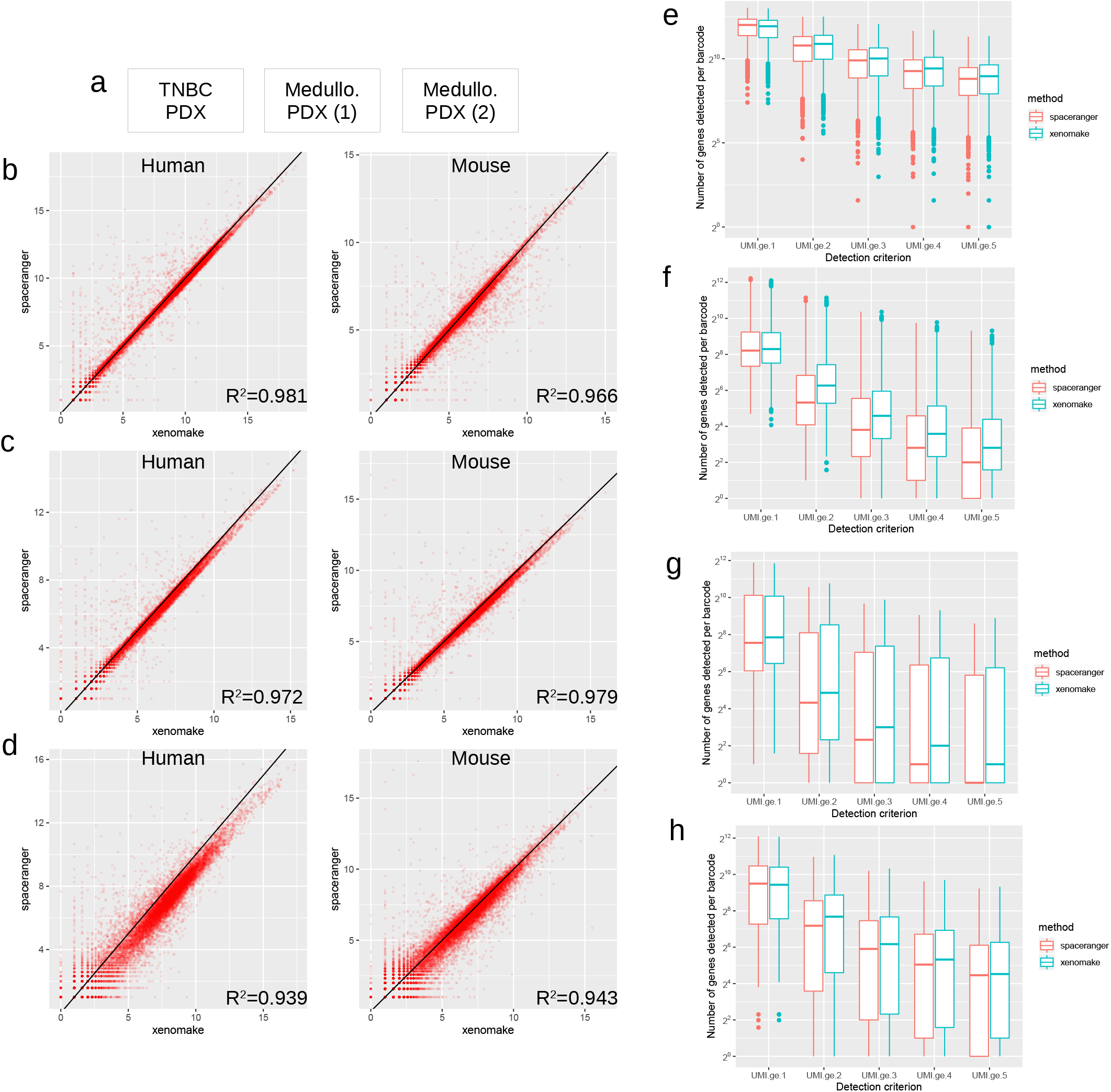
Comparison between Xenomake and Space Ranger. **a**. Datasets used. **b-d**. Correlation in total UMI count per gene between the two methods. Each dot: a gene. Line: identity. **b**. TNBC PDX data comparison in human (left) and mouse (right) matrices. **c**. Medulloblastoma PDX (palbociclib) data comparison in human (left) and mouse (right). **d**. Medulloblastoma PDX (control) data comparison. **e-h**. Number of genes detected per spatial barcode for various detection thresholds (UMI count>=1, 2, 3, 4, 5). UMI.ge.1 stands for UMI greater or equal to 1. **e**. TNBC PDX human portion. **f**. TNBC PDX mouse portion. **g**. Medulloblastoma PDX (palbo) human. **h**. Medulloblastoma PDX (palbo) mouse.

We compared our tool with Space Ranger where reads were aligned with the mouse-human integrated genome. Our comparison focused on the genic reads among the in-tissue spatial barcodes judged by Space Ranger [6]. Xenomake mapped a total of 17,126 human genes, and 14,647 mouse genes among 2,217 in-tissue barcodes. Compared to Space Ranger, the total aligned human genic reads in these barcodes increased from 29.5 million to 32 million, while the increase on the mouse side was from 2 million to 2.2 million reads. The correlation in total UMI count per gene between the two methods is very high (R2>0.98), but Xenomake is able to assign more reads per gene than Space Ranger (**Figure 2b**). The majority of genes are distributed below the identity line towards Xenomake in the correlation plot. Xenomake also significantly improves the number of genes detected per spatial barcode (**Figure 2e-f**). At each detection threshold of UMI count>=2, 3, 4, 5, the genes detected per barcode are greater in Xenomake than Space Ranger, with the differences more prominently exhibited in mouse. (**Figure 2e-f**).

Besides our own dataset, we also applied Xenomake on a recent ST dataset focused on medulloblastoma PDX samples in the control and palbociclib treated setting (**Figure 2a**). We similarly observed an increase in the number of genes detected per barcode at each detection criterion of UMI count>=2, 3, 4, and 5 (**Figure 2g-h**). The vast majority of genes have a higher total read count (comparing Xenomake with Space Ranger) (**Figure 2c-d**). These results suggest that Xenomake reduces the sparsity of the gene expression matrix by increasing read counts on a per gene and per spot level.

Finally, we compared UMI counts in the gene-by-barcode output matrix to count the number of entries where (1) the counts between Xenomake (X) and Space Ranger (S) remained the same, (2) S>X, and (3) X>S (**Figure 3**). We compared only entries which are non-zero in either X or S. The number of cases of X>S range from 23% to 59% of total non-zero entries (representing improvements made by Xenomake) for the 3 datasets (**Figure 3**), and are higher than S>X.

**Figure 3:**
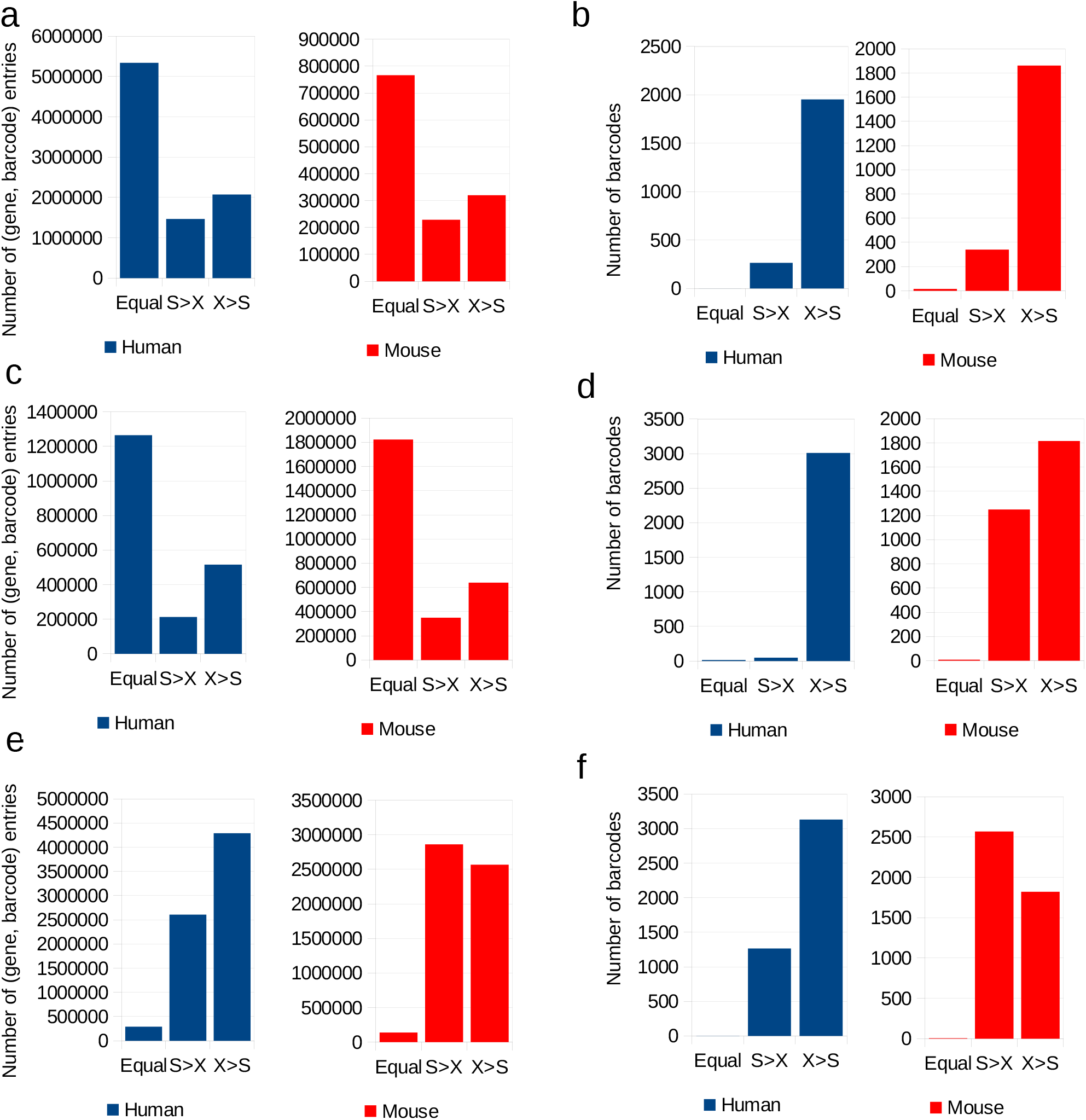
Comparison between Xenomake and Space Ranger on a per (gene, spatial barcode) basis. **a, c, e**. We utilized all in-tissue barcodes and genes with nonzero total UMI count. Out of all nonzero entries in the gene-by-barcode matrix, we count how many entries is Xenomake (X) count equal to Space Ranger count (S), greater than (X>S), or less than (S>X) Space Ranger. **a-b**. TNBC dataset. **c-d**. Medulloblastoma (palbociclib) dataset. **e-f**. Medulloblastoma (control) dataset. **b, d, f**. Comparison in terms of total UMI counts per barcode. Barcodes are similarly put into three categories: Equal, X>S, or S>X.

### Xenomake finds stroma-specific genes and secreted cytokines

To determine if Xenomake preserves biological context, we next checked the expression of stromal cell type specific genes in the mouse gene expression matrix (**Figure 4**), as we expected that stromal expression should be low in human but high in mouse. Conversely, epithelial expression should be high in human and low in mouse transcriptomes. Expectedly, in the TNBC PDX dataset, Pecam1, Fcgr3, Csf1r expression, which respectively mark endothelial cells, NK/neutrophils, and macrophages [24–26], are high in the mouse portion, but lowly expressed in the human counterparts (PECAM1, FCGR3A, CSF1R) (**Figure 4a**). Additionally, collagen Col1a1, an abundant protein in the extracellular matrix (ECM), is much higher than COL1A1 in human (**Figure 4a**). These results suggest that our tool Xenomake can correctly assign many stromal exclusive transcripts to the mouse transcriptomes.

**Figure 4:**
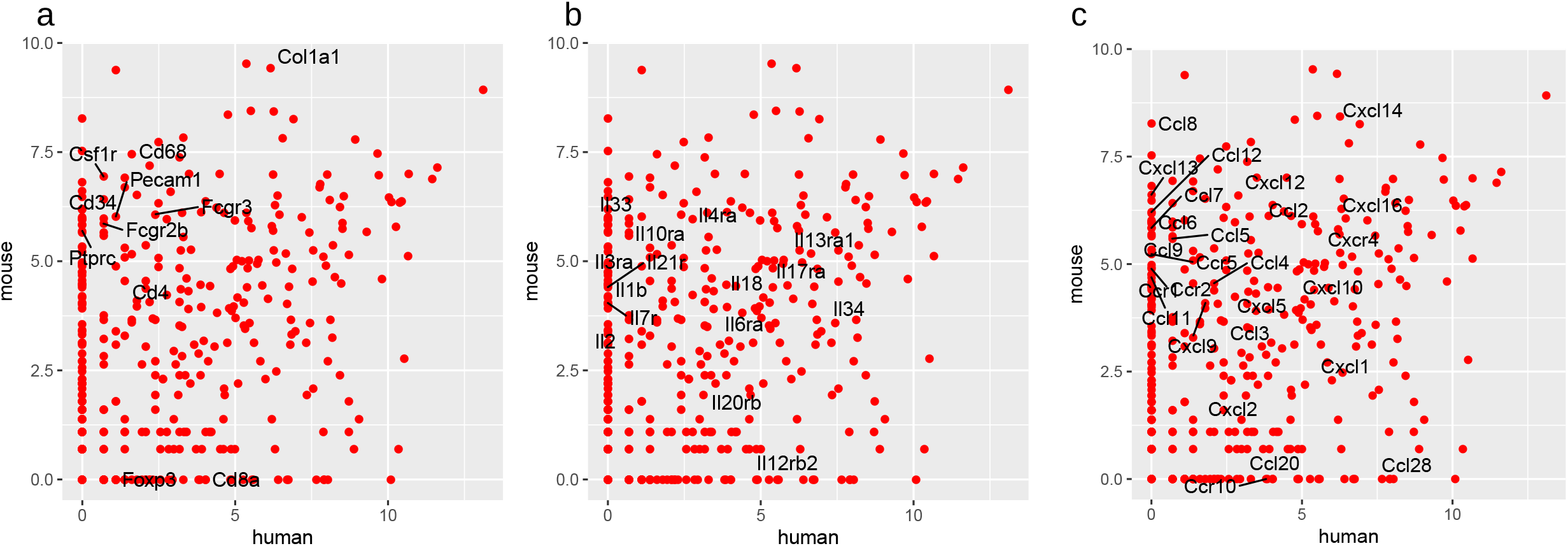
Human/mouse homolog expression in TNBC, illustrating differential cell type marker, interleukin, and cytokine expression. Each dot plots the mouse and human homolog expression after PDX reads are partitioned to each organism. For example, the dot marked Col1a1 indicates the expression Col1a1 in mouse plotted against the expression of COL1A1 in human. For fully labeled scatterplot, please refer to Supplementary Fig 3. **a**. Stromal cell-type specific genes which display mouse bias. **b**. Interleukin genes. **c**. Cytokine and chemokine genes.

Chemokines and cytokines are simultaneously secreted by both stromal cell types and epithelial tumor cells in the tumor’s microenvironment and are an integral part of cell-cell signaling [27,28]. It is often difficult to attribute chemokines expression to stromal or tumor cells in standard models, but in PDX models by dividing transcriptomes into human and mouse, one can study the differential production of chemokines and identify biomarkers in stromal and epithelial compartments [29]. We therefore checked the expression for interleukins (**Figure 4b**), cytokine/cytokine receptor, and chemokine/chemokine receptor pairs (**Figure 4c**) for evidence of ligand-receptor pairs displaying stroma and epithelial bias. We found CCL28–CCR10 to be primarily human biased in the TNBC model (**Figure 4c**), suggesting that these chemokines are key to epithelial cells communication with the surrounding environment. In stroma, interleukin Il33 and Il10ra are upregulated in stroma compared to epithelial cells (**Figure 4b**). Il33 and its receptor Il1r1, which also is stroma biased, is known to contribute to Th2 and regulatory T cell (Treg) functions [30]. Il33 induces inflammatory response through NFKB signaling [31]. Thus, we suggest that Xenomake can accurately identify differential cytokine gene expression in PDX models and will enable researchers to investigate cell signaling in tumor microenvironments.

### Discrepant genes between Xenomake and Space Ranger

We next selected the top 50 genes that are ranked high in Space Ranger but low in Xenomake output, and 50 genes in the reverse direction (**Supplementary Fig 1**) in the TNBC dataset. We checked for evidence of expression for these genes in a separate single-cell RNAseq breast cancer atlas (**Figure 5a, b**). After filtering for overlap with scRNAseq data, we arrived at 41 genes. In the mouse compartment showing Xenomake-high genes (**Figure 5a**), 17 genes are enriched for stroma cell-type specific expression in endothelial cells and myeloid cells in the scRNAseq atlas (see red boxes **Figure 5a**). This list encompasses important transcriptional regulators and RNA processing genes such as Satb1, Bcl11a, Foxc1, Tox, Ptbp2, and Hnrnpd (**Supplementary Fig 1a**). In contrast, in the mouse compartment, many Space Ranger-high genes are also expressed highly in cancer epithelial cells (see blue box **Figure 5b**), suggesting possible non-specificity of some of Space Ranger output. We quantified the cell-type specificity of Space Ranger-high and Xenomake-high genes using Shannon Entropy. Xenomake generally has a lower entropy score, meaning higher specificity, than Space Ranger (**Figure 5c**), particularly for endothelial, myeloid, PVL, and T/B cells (**Figure 5d**). In the human compartment, we found over 30 out of 46 discrepant Space Ranger-high genes to lack cancer cell expression in the scRNAseq atlas (**Supplementary Fig 2**). Overall, these results indicate that where results between these two methods are discrepant, Xenomake’s results are more likely to find extra support from external scRNAseq data.

**Figure 5:**
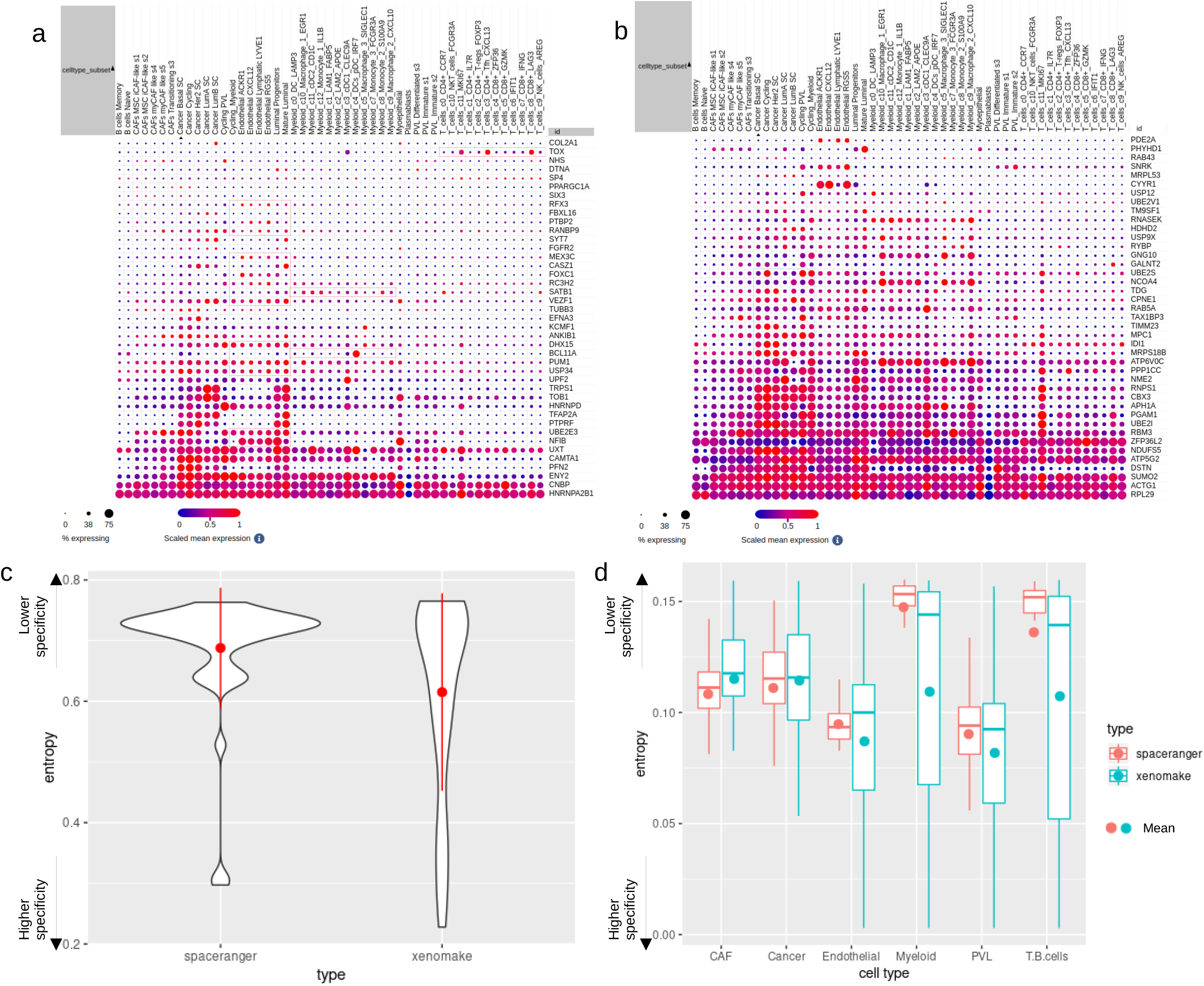
Xenomake- and Space Ranger-biased genes in the mouse compartment and their evaluation using scRNAseq (TNBC). Genes that were high in Xenomake and low in Space Ranger, and vice versa, were extracted from the mouse matrix (Supplementary Fig 1a). The expression levels of these discrepant genes were next visualized using an external breast cancer single-cell atlas (**a-b**). **a**. Xenomake-biased genes. Red boxes: genes showing evidence of stroma cell-type expression according to scRNAseq. **b**. Space Ranger-biased genes. Blue boxes: genes showing non-specificity in epithelial cancer cells. **c**. Stroma cell-type specificity scores, measured in entropy. Lower means better. Red dot and vertical bar: mean +/- 1 standard deviation. **d**. Entropy divided by cell-types. Dot: mean.

## Discussion

As spatial transcriptomics becomes utilized more and integrated with PDX models, the need for a reproducible workflow to handle these projects becomes increasingly necessary. We have introduced Xenomake, which is an end-to-end pipeline that includes read alignment and gene quantification steps for reads generated by spatial transcriptomic platforms. We provide tools to confidently assign reads to different genomes used in PDX models and methods that recover more reads, maintain robustness of datasets, and preserve biological significance.

Xenomake is written using Snakemake [15], a workflow management system that is designed to increase reproducibility for analyses. Snakemake offers a framework for projects so that they are human readable, customizable, and scalable. We believe that by implementing our pipeline with Snakemake, we offer reproducibility in future spatial PDX projects and provide a framework for subsequent spatial pipelines to build upon. Xenomake is an independent and standalone software that does not require Cell Ranger [33] or Space Ranger [6] to run. As such, it is an appealing open-source alternative to Space Ranger, and its open-source license encourages programmatic development from the community. Of note, Space Ranger does not currently utilize a specialized xenograft sorting program for PDX reads. We showed that Xenomake is superior to standard methods in handling spatial xenograft samples in a number of measures.

A key issue with current methods is the loss of potentially biologically significant reads. In PDX datasets, mouse reads can already be sparse due to low stromal infiltration in some cancers. We show that we can recover more reads compared to standard workflows, by more intelligently utilizing “both” and “ambiguous” categories in Xenograft/Xenome output. Higher read coverage will likely translate to increased power and interpretability of downstream analyses. Currently, Xenomake only handles Visium [12] spatial transcriptomics experiments. However, we believe it will be important moving forward to extend this framework to handle other spatial and single-cell datasets such as Slide-Seq [13] and scRNA-seq [32]. Providing an extendable framework to handle all possible PDX projects is important to further anticipate and meet future needs.

## Methods

### PIM001P PDX Animal Study

PIM001P was obtained from the University of Texas MD Anderson Cancer Center where it was generated and characterized through a materials transfer agreement. PIM001P was propagated in mice as previously described [34]. In brief, frozen PDX cell suspensions were thawed and washed with Dulbecco’s modified Eagle’s medium (DMEM):F12 (Cytiva HyClone, SH30023.01) supplemented with 5% FBS. Cells were stained with AOPI (Nexcelom Bioscience, CS2-0106) to count viable cells on a Cellometer K2 (Nexcelom Bioscience). 0.5 million viable tumor cells were suspended at a 1:1 ratio in Matrigel (Corning, 354234). Suspended cells were then injected into the fourth mammary fat pads of NOD/SCID mice [NOD.CB17-Prkdc^scid^/NcrCrl, Charles River, National Cancer Institute (NCI) Colony] aged 5 weeks old.

The tumor was harvested 2 days after it obtained a volume of 150mm3. At time of harvest, a thin slice of fresh tumor was cut and placed in a cassette. The cassette was then filled with O.C.T. (Sakura Finetek, Catalogue 4583) and submerged in isopentane until frozen solid. Frozen tissue cassettes were stored at -80C.

### Library Preparation and Sequencing of PIM001P

Tissue sections of 10µm thickness were mounted onto the capture areas of the Visium Spatial Gene Expression slide and stained using hematoxylin and eosin. Tissue sections were permeabilized on a thermocycler for 24 minutes, as determined by the Tissue Optimization step. Poly-adenlyated mRNA is released and captured by surface-bound primers within each capture area. Reverse transcription, template switching, extension, and second strand synthesis are performed on the slide. Full-length, spatially barcoded cDNA transcripts are then denatured from the slide and amplified via PCR prior to library construction. Approximately 110 to 375 ng of amplified cDNA was carried forward into library construction. During library construction, cDNA is enzymatically fragmented to target amplicon size then undergoes end repair, A-tailing, adapter ligation, and then amplified using between 14 and 16 PCR cycles. The resulting libraries were quantitated using the Invitrogen Qubit 2.0 quantitation assay and fragment size assessed with the Agilent Bioanalyzer. A qPCR quantitation was performed on the libraries to determine the concentration of adapter ligated fragments using Applied Biosystems ViiA7 Real-Time PCR System and a KAPA Library Quant Kit (p/n KK4824). All samples were pooled equimolarly and re-quantitated by qPCR, and also re-assessed on the Bioanalyzer.

### Sequencing

150 pM of equimolarly pooled library was loaded onto the NovaSeq 6000 S4 flowcell and sequenced at the recommended 28-10-10-50 read configuration. PhiX Control v3 adapter-ligated library (Illumina p/n FC-110-3001) was spiked-in at 2% by weight to ensure balanced diversity and to monitor clustering and sequencing performance. A minimum of 300 million read pairs per sample was sequenced. FastQ file generation was executed using 10X Genomics’ Space Ranger mkfastq software.

### Xenomake Pipeline

#### Input

Our pipeline requires paired end FastQ files as input for our pipeline and we recommend using Space Ranger [6] mkfastq to make this conversion from raw Illumina base calls. We utilize Picard’s [18] “FastqToSam” to convert FastQ files to an unaligned BAM file. This format allows preprocessing such as tagging and trimming of reads prior to alignment.

#### Transcript Tagging

Unique molecular identifiers (UMIs) and cell barcode information is present in predictable locations of forward reads as part of illumina chemistry. This positional information can be utilized to extract read segments and tag both paired end reads. For example, UMIs are always found in positions 17-28 and cell barcode information is found at positions 1-16 on forward reads.Drop-seq [17] provides the function TagBamWithReadSequenceExtended to extract bases and create a new BAM tag with those bases on the genome read. For cell barcode tagging, we use the following parameters; “BASE_RANGE=1-16 TAG_NAME=CB”. For UMI tagging we use “BASE_RANGE=17-28 TAG_NAME=MI”

#### Adapter and PolyA Trimming

The Drop-seq [17] program provides tools to remove adapter sequences from 3’ ends of reads and polyA stretches using TrimStartingSequence and PolyATrimmer respectively. PolyATrimmer is designed to trim polyA tails from reads by searching for at least 6 contiguous bases at the 3’ end with zero mismatches and hard clips of these bases (MISMATCHES=0, NUM_BASES=6) Additionally TrimStartingSequence searches for 5 contiguous bases with zero mismatches in a provided Illumina adapter sequence (SEQUENCE=Adapter_Sequence MISMATCHES=0 NUM_BASES=5) and hard-clip the bases off the read if they occur at the 5’ end.

#### Mapping

Mapping is performed via STAR [7] on tagged and trimmed unaligned bam files. We utilized default mapping and indexing parameters for genome assemblies provided by STAR and output an unsorted bam file. Since STAR doesn’t preserve cell barcode and UMI tags, these need to be merged from the unaligned bam file to the aligned bam output. A custom python script is used to merge tags to generate a tagged and aligned bam file. Subsequently, the Drop-seq [17] program TagReadWithGeneFunction is used with default parameters to tag reads that are exonic when the read overlaps the exon of a gene. This tag contains the name of the gene as reported in the provided annotation file and a tag from the following list [INTERGENIC, INTRONIC, UTR, CODING, RIBOSOMAL].

#### Xenograft Sorting

The core of Xenomake is the implementation of xenograft sorting on reads that are mapped to both mouse and human organisms (overlapped reads). Xenograft Sorting is performed using Xengsort [11], a faster and lightweight implementation than previous methods such as Xenome [10]. Xengsort is an alignment-free method that stores k-mers into a large table that enables the lookup of species of origin (host,graft, or both) of each RNA k-mer that occurs in either genome assembly. Reads are classified by iteratively searching k-mers for the appropriate species of origin. This process assigns reads into “host”, “graft”, “ambiguous”, “both”, or “neither” based upon Xengsort’s decision rule. When building the index files used in classification, Xengsort recommends storing 4.5 billion 25-mers when using humans and mouse genomes. After index generation, we implement Xengsort classify on our reads using default settings.

#### Reassignment of Xenograft Sorted Reads

We can confidently reassign reads back into the human or mouse alignment files on xenograft sorted files (graft and host respectively). Our idea for handling multimapped reads is inspired by another spatial processing tool, Spacemake [35]. Briefly, we wrote a Python script that can assign the “ambiguous” and “both” output reads back into respective organisms. It compares gene function tags and STAR alignment scores (AS) to determine which organism this read should be reassigned. Briefly, a read is compared between both organism’s alignment files side by side and makes a call based upon the following strategies. First, if only one of the reads aligns to an exonic region, then it is reassigned to that organism. Second, if the AS differs between two organisms, the read with a higher AS is reassigned. If neither requirement is satisfied, the read is discarded. This method leverages information from mapping, functional tags, and alignment free sorting to recover as many reads as possible.

#### Within-species Multi-Mapped Read Assignment

Handling multi-mapped reads within species works by filtering reads that are multi-mapped to different genomic positions by STAR [7]. Its decision rule is identical to the reassignment of xenograft reads but does no organismal comparison. Again, it prioritizes reads that are mapped to exons, have a higher alignment score, and removes the “worse” read from the alignment. This is an optional parameter but removes the potential to count reads twice in downstream processing.

#### Output Files

We offer multiple options for files used in downstream analysis. First, a gene by spot umi count matrix is generated using the Drop-seq [17] program DigitalGeneExpression. This takes the gene tags, cell barcode tags, and UMI tags to develop a count matrix in tab delimited format. We also implement a custom script that replicates the 10X genomics HDF5 architecture. Tools such as Scanpy [20], Giotto [22], and other spatial pre-processing can take HDF5 spatial files as input and allow easier integration for downstream analysis by users. Additionally, we also utilize Scanpy to generate H5ad objects that can be loaded directly by multiple spatial packages.

#### Downstream Processing

We utilize the H5ad object and a Scanpy workflow [36] to provide basic downstream analysis of spatial samples as an optional parameter. As a part of this, we offer an estimation for in-tissue segments in-silico using KMeans clustering. We set the number of clusters a priori to two and grouped all spots into these categories. Fundamentally, spots with low expression and low number of genes expressed should group together; conversely spots with high counts and higher number of genes should cluster and represent in-tissue segments.

We further process samples by filtering spots that are designated out-of-tissue. We subsequently find highly variable genes (HVGs) using sc.pp.highly_variable_genes and use a subset of HVGs to perform principal component analysis (sc.pl.pca), UMAP projection on the top 40 PCs (sc.pl.umap) and perform Leiden clustering with default parameters (sc.pl.leiden). The output is saved as an Anndata object and can be loaded with spatial transcriptomics packages such as Scanpy and Squidpy.

### Analyses

#### Comparison with Space Ranger

We downloaded Space Ranger version 2.0.1 from the 10X Genomics website. Following the recommended tutorial, we built a reference for multiple species by running the “spaceranger mkref” command with the two reference genome assemblies being mm10 (version M23 Ensembl 98) and GRCh38 (version 32 Ensembl 98). We next used “spaceranger count” with default settings to align reads to the integrated reference genome and quantify gene expression. The output of Space Ranger attached “hg38_” and “mm10_” prefixes to each human and mouse gene name, allowing further division into individual species. The capture area information, i.e., A1-C1, was available from the Visium slide for TNBC PDX and on the raw data repository for the medulloblastoma PDX. Following quantification, we extracted in-tissue barcodes and all genes with nonzero read counts in these barcodes to use them for comparison purpose. For per-gene and per-barcode comparisons, we summed gene expression across in-tissue barcodes for each gene, and summed gene expression across all genes for each barcode. For the number of genes detected per barcode, we set a gene expression detection threshold criterion to 2-5 UMI counts and counted the number of genes exceeding the detection threshold. When making comparisons between methods, we counted the number of cases in the three categories: (1) Equal: Xenomake and Space Ranger gene counts are the same, (2) X>S: Xenomake returns higher gene counts than Space Ranger, and (3) S>X. Intuitively, (1) will indicate agreement between methods, while (2) relative to (3) will indicate the superiority of Xenomake in terms of increased read depth.

#### Stroma analysis

We extracted all interleukin-related and cytokine-related genes using a regular expression tool that detects “IL”, “CCL”, “CCR”, “CXCL”, and “CXCR” in gene names. Then we checked each gene to determine if it has a homolog in mouse and removed those that did not have a mouse homolog. We deduced a list of human-mouse homologous pairs (e.g. CCL10—Ccl10). We plotted the expression of human homolog against the mouse homolog. Those homologs that exhibit +/- 2 standard deviations from the diagonal line in the scatter plot were deemed to have an epithelial or stromal bias.

#### Comparison using single-cell RNAseq

We downloaded a breast cancer single cell atlas from Wu et al. Cell type specific gene expression was visualized using a dot plot. To extract discrepant genes between Xenomake and Space Ranger, we determine the top 50 genes, in each direction, that lie furthest from the diagonal line in the correlation plot (Figure 2). These genes were labeled in Supplementary Figure 1. To compute cell type specificity score for a gene *g*, we utilized the Shannon Entropy Index which is defined as:

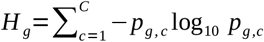

Where *H*_*g*_ is the entropy of *g, c* is a cell-type, *p*_*g,c*_ is the probability of expression of *g* in cell-type *c*, computed as the sum of expression of *g* in all single cells in cell-type *c* divided by the sum of expression of *g* in all cells.

## Acknowledgements

This project was supported in part by the Genomic and RNA Profiling Core at Baylor College of Medicine with funding from the NIH S10 (1S10OD023469) and NIH NCI (P30CA125123) grants. QZ is supported by Cancer Prevention and Research Institute of Texas (CPRIT) grant RR220035. GVE is supported by grants RR200009 from CPRIT, R37 CA269783-01A1 from National Cancer Institute (NCI), and RSG-22-093-01-CCB from American Cancer Society (ACS). KEP is supported by F31CA275397 from NCI. The generation of PIM001-P was supported by a generous gift from the Cazalot family and the MD Anderson Women’s Cancer Moonshot Program. GVE is a CPRIT Scholar in Cancer Research. QZ is a CPRIT Scholar in Cancer Research. Computing was done using the Dan L Duncan Cancer Center (DLDCC) cluster at BCM (Baylor College of Medicine) with 30 heterogeneous nodes, each with 24-64 cores and up to 256 GB RAM, running Red Hat Enterprise Linux with queues managed by SLURM. We thank Michael Dehart for providing computing support.

## Availability of data and materials

Xenomake is maintained on the Github repository located at https://github.com/istrope/Xenomake/. Installation of Xenomake requires a Conda environment and Snakemake. More details can be found at the repository website. Medulloblastoma PDX data can be accessed at the accession ID E-MTAB-11720 on ArrayExpress (https://www.ebi.ac.uk/biostudies/arrayexpress/studies/E-MTAB-11720). Raw data for TNBC PDX generated in this study has been deposited on Zenodo https://zenodo.org/record/8313189.

## List of Figures and Supplementary Figures

**Supplementary Figure 1: a**. Full list of Xenomake-biased and Space Ranger-biased genes in the mouse compartment. Correlation plot is the same as Figure 2c. **b**. Biased genes in the human compartment. Correlation plot is the same as Figure 2b.

**Supplementary Figure 2:** The expression levels of discrepant genes found from the human compartment in the breast cancer single-cell RNAseq atlas. **a**. Space Ranger biased genes. Blue boxes: genes that do not have appreciable expression. **b**. Xenomake biased genes.

**Supplementary Figure 3:** Full list of human/mouse homolog pairs.

