## Supplementary figures and images for "Xenomake: a pipeline for processing and sorting xenograft reads from spatial transcriptomic experiments"

### Supplementary Figure 1

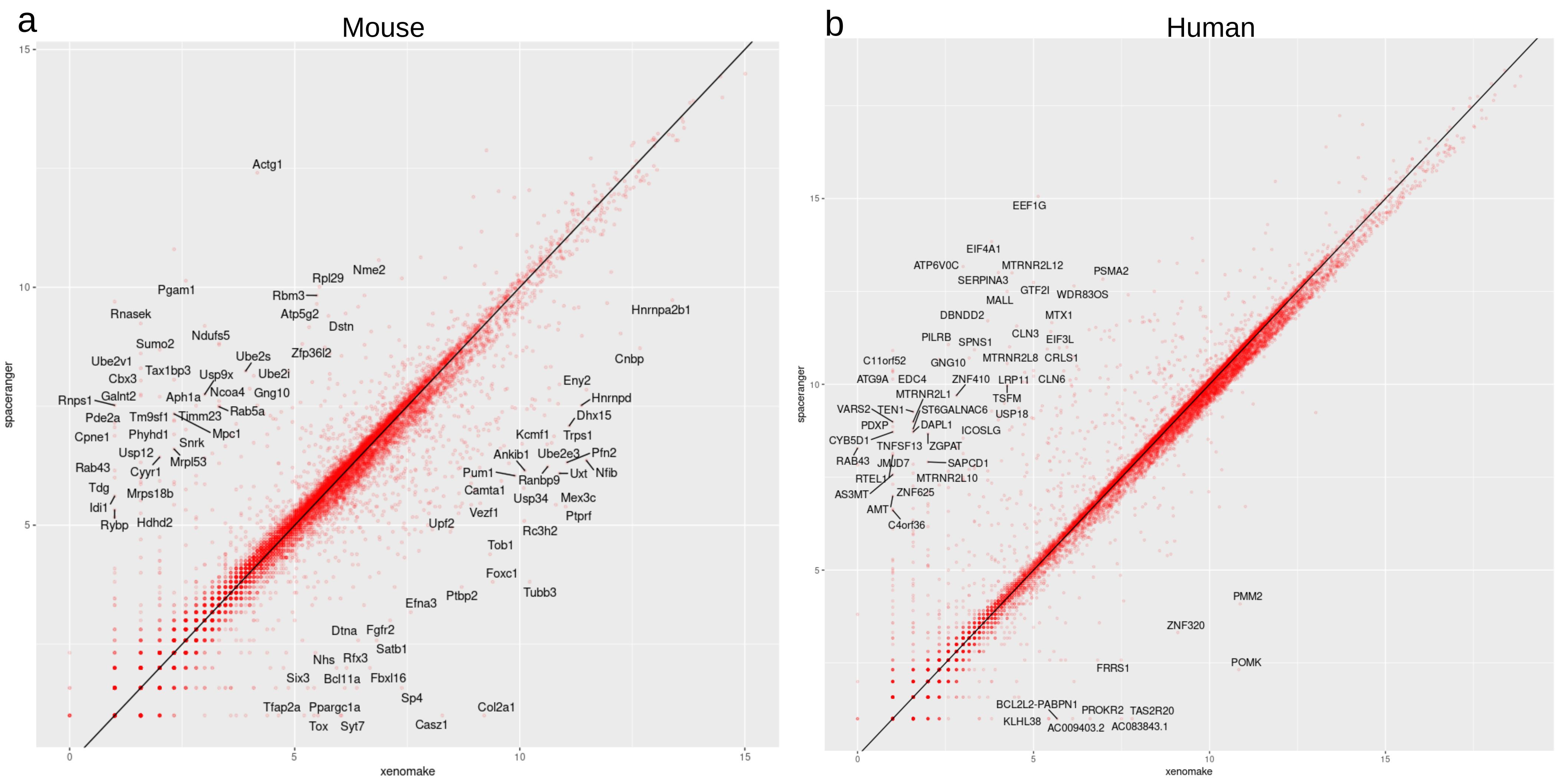

### Supplementary Figure 2

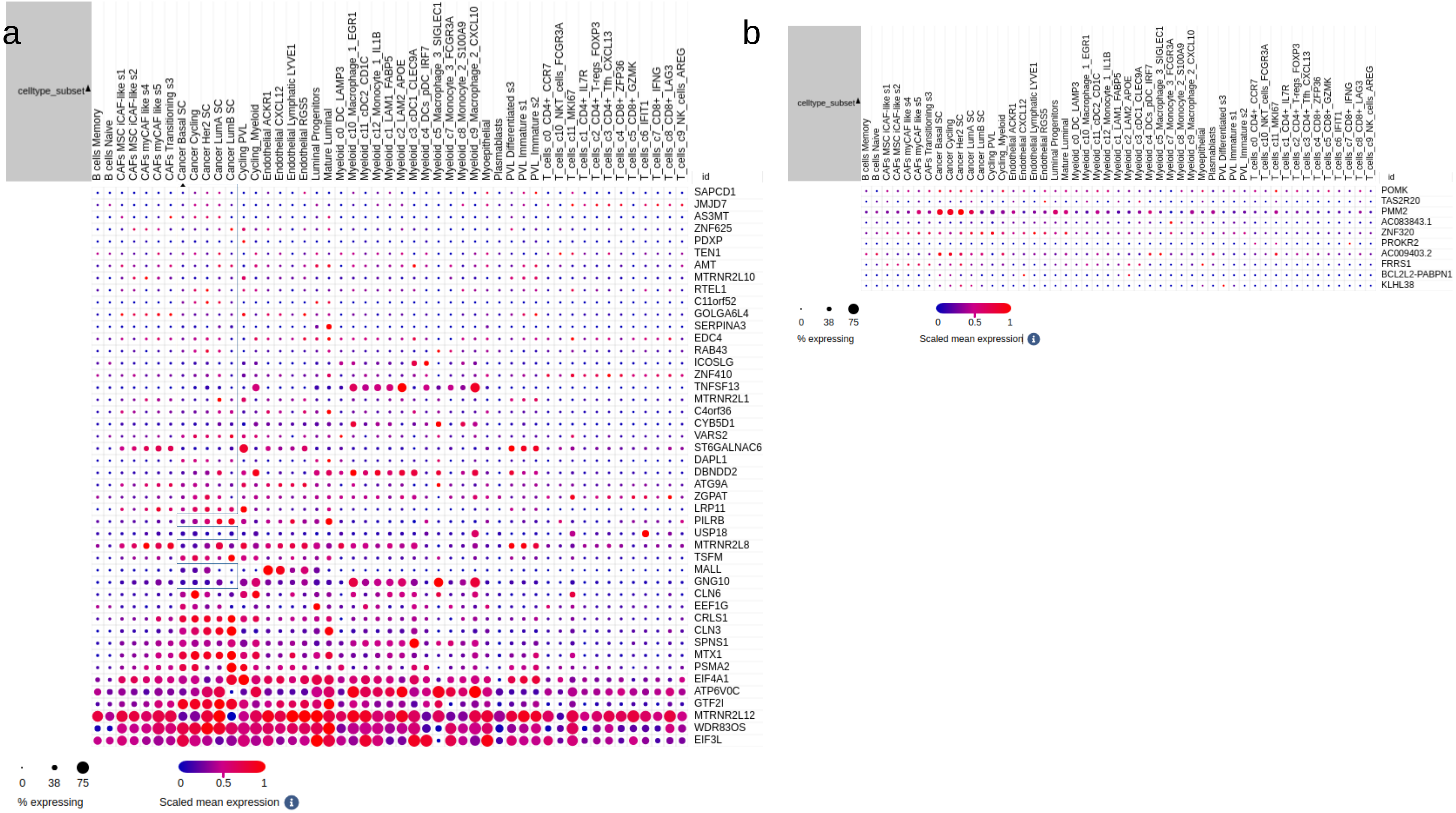

### Supplementary Figure 3

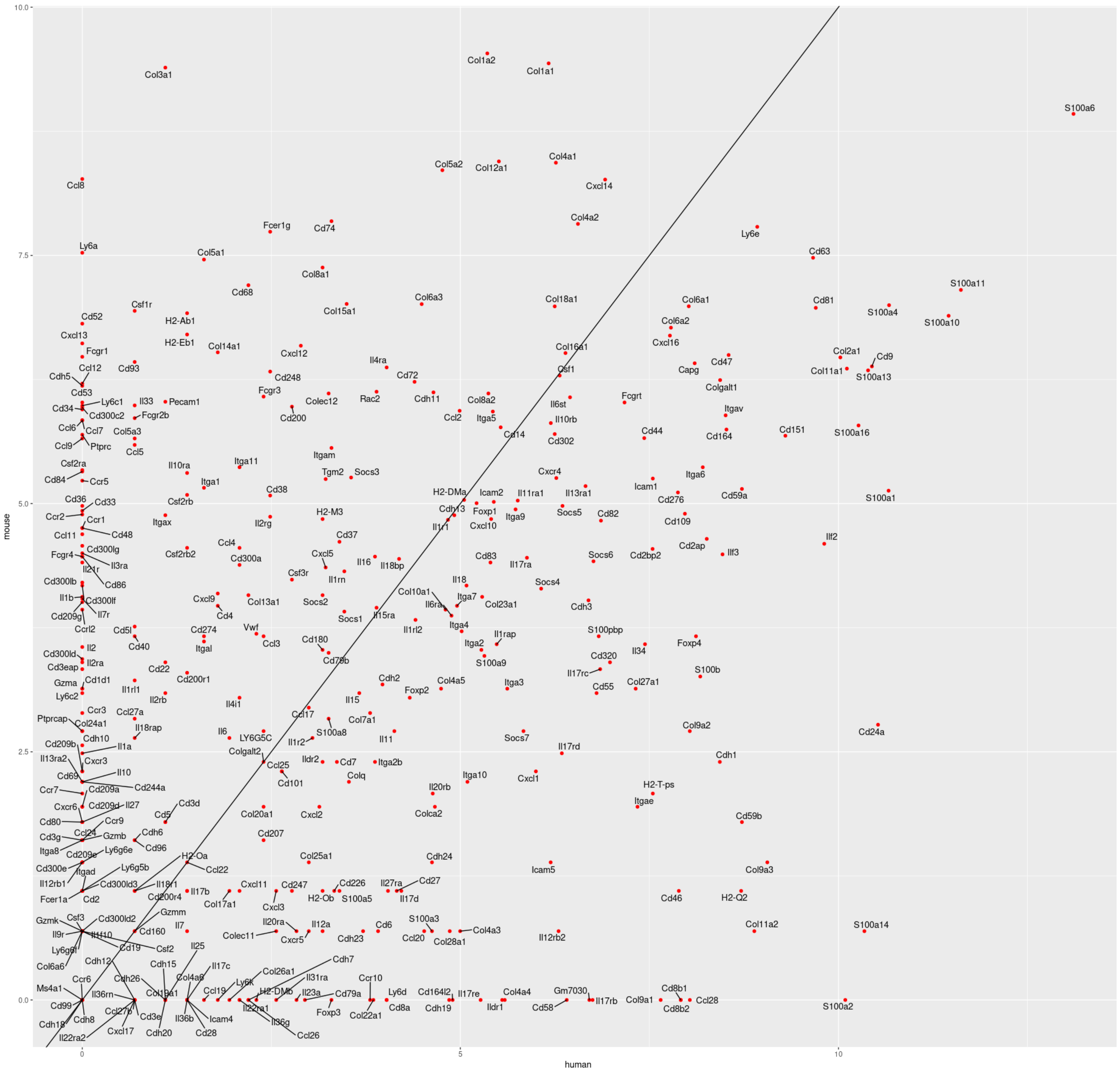
